# Proliferation and mutation as humoral immune response

**DOI:** 10.64898/2026.11.06.764896

**Authors:** Isaac Meilijson, Laura Lea Sacerdote

## Abstract

The affinity maturation process, through which antibodies acquire relatively high affinity for an invading pathogen, involves complex mechanisms whose relative importance is not yet fully understood. This study proposes an analytical model of this process, to investigate the main factors governing proliferation and mutation and assess their respective roles. The model suggests that the immune system should not be expected to follow a single, rigid evolutionary pattern, as a wide range of proliferation and mutation strategies can yield nearly optimal outcomes.

## 1 Introduction

The ability of the immune system to recognize an invading pathogen and to produce a suitable antibody is crucial for a species’ survival. In the so-called affinity maturation step, activated B-cells proliferate and mutate, and the best ones are selected to differentiate into plasma cells, that secrete high-affinity antibodies to attack the pathogen. Repetition of this cycle increases the binding strength, i.e., the affinity for the pathogen, by enhancing antibody specificity toward the target over time.

This study applies a mathematical model to describe the affinity-growth mechanism for antibodies that recognize an invading pathogen with a relatively high affinity and then start the B-cells proliferation/mutation cycle to further increase it. The aim is twofold: on the one hand, to recognize the main proliferation and mutation factors that determine efficient affinity growth; on the other, to investigate the flexibility in the determination of these factors to achieve a small probability that the pathogen kills the host.

Similarly to Agur et al. (1991/2) and Merkenschlager et al. (2025), the analytical model adopted in this study assumes that the immune system tools available for creating antibodies with higher affinity are an accelerated rate *λ* of B-cell division (or proliferation) and accelerated rate *ρ* of base pair (or point) mutations along the DNA sequence encoding the variable region of the antibody. The pathogen has the ability of killing the host with lethality rate *r* = *Rλ*, where the relative lethality parameter *R* should be thought of to be for illustration purposes between 0.0001 and 0.01. *R* will be assigned throughout the three values 0.01, 0.001 and 0.0001. The rate *λ* is commonly quoted as reflecting division of a B-cell in 8 to 10 hours on the average. An unrecognized pathogen will kill the host after 37 days on the average if *R* = 0.01 and after 10 years if *R* = 0.0001.

Agur et al. worked under the postulate that one favorable mutation is enough to create the high affinity antibodies capable of eliminating the pathogen. The current study relaxes this assumption and investigates whether high affinity can still be achieved when the immune system involves successive cycles that progressively increase antibody affinity. Under the proposed analytical model of the immune system, each cycle undergoes a *basic* strategy with some parameters *n* and *ρ*, which consists of pure B-cell proliferation until a clone of size *n* has evolved, followed (in parallel to further B-cell proliferation) by point mutation at a fixed mutation rate *ρ* until a B-cell with antibodies of higher affinity has been created. The *global* strategy, or **proliferation/mutation management**, consists of the repeated application of basic strategies until the emergence of an antibody that can eliminate the pathogen. The host survives if this entire process is concluded before the pathogen kills the host.

### Optimal, standard and bang-bang proliferation/mutation management

Management strategies could in principle change the mutation rate gradually as the clone size changes. Agur et al. showed (in their setup) that the optimum is the biologically infeasible “bang-bang” management where mutation rate is zero until a threshold clone size has been reached, then infinite for a split moment, and back to zero if the single mutant was adverse, waiting for the next B-cell proliferation. The best basic (*n, ρ*) strategy where mutation rate is zero until reaching the threshold clone size *n*, and then *ρ* is kept constant until a favorite mutation emerges, was shown to yield practically the same survival probability as the best bang-bang management. This is the rationale for resorting throughout this study to the basic and global strategies based on fixed mutation rates onset by threshold clone sizes. The term “optimal” will be applied to the best such management, and the term “standard” will be applied to management strategies where the same pair (*n, ρ*) is applied at every affinity level and against every risk *R*.

### Diversity of efficient strategies

A salient result of the analysis to be conducted through the mathematical model is that the survival probabilities under the two types of management are indeed similar: the survival probability cannot be significantly increased by distance-and-risk-dependent management. The host does not need to know in advance the risk posed by the current pathogen, a request that is not biologically realistic, nor to react differently depending on the degree of affinity. Due to the implied high variety of nearly optimal management strategies, the immunological response to a pathogen, often referred to as “not well understood”, need not follow a predictable pattern. On the other hand, as is commonly accepted, a high variety of nearly optimal management fosters evolution efficacy.

The basic strategy re-start nature of the global strategy postulates, in particular

### Working assumption

Once a higher affinity antibody has been created, the clone of B-cells that generated it ceases to be active.

This assumption does not restrict efficiency in a critical way: as will be shown by simulation. If the lower affinity B-cells continue to supply a steady stream of mutating antibodies, the effect will be a relatively minor proportional increase in the survival probability. However, this increase comes at the cost of magnifying 5 to 10-fold the trail of adverse mutations.

### Affinity and distance

In the quantitative language of “distance” trying to accomodate the biological notions of “affinity” and “specificity”, the original antibody recognizes the antigen at some distance *k*, and the basic strategy concludes with the creation of a B-cell with antibodies at distance *k* −1. The global strategy is successful if an antibody at distance *k* = 0 is created before the pathogen kills the host. The DNA length encoding the variable binding region of the antibody with the antigen (henceforth, DNA length) is denoted by *m*, generally postulated in the recent literature to be between 200 and 260 (of which *k* is a single digit), by adding together the two chains VH and VL (heavy and light). The distance *k* represents the number of favorable mutations the B-cell has to undergo to create an antibody that can eliminate the pathogen. The current study considers the two chains as one, and assumes that point hypermutation chooses to mutate a randomly chosen amino-acid among the *m*. At the considered level of detail, no differentiation is made between framework (FR) and complementarity-determining (CDR) subregions of the binding region (see Zhu et al. (2024)), and the degree of affinity is quantified solely by distance, ignoring other important measures like antibody concentration.

In the immunological system some favorable mutations may achieve higher affinity than others, and “distance” may be a scenario-dependent process. The adopted mathematical model postulates the rigid linear pattern that the initial distance is some scenario-dependent *k*, but each favorable mutation reduces the distance by 1. Following the literature that sets as 1 to 5 and above the number of mutations to achieve a B-cell that secretes high-affinity antibodies that can attack the pathogen, the range of distances *k* to be analyzed is from *k* = 1 to *k* = 6.

Recognition at distance *k* = 1 achieves higher survival probability than recognition at lesser affinity. In contrast, as the potential repertoire of antibodies close to the pathogen’s antigen increases very fast with *k*, much fewer “scouts” are needed if *k >* 1. Furthermore, if a B-cell with antibodies at distance *k* = 0 was trained in a previous invasion, recognition at higher distance is an effective response to minor antigen mutations undergone by the pathogen.

### Proliferation/mutation strategies

A global strategy (*n*_1_, *ρ*_1_), (*n*_2_, *ρ*_2_), …, (*n*_*k*_, *ρ*_*k*_) is a sequence of basic strategies. If the current B-cell with closest antibody to the pathogen’s antigen is at distance *i* (1 ≤ *i* ≤ *k*), apply pure proliferation until a clone of size *n*_*i*_ of this B-cell has been created, followed (in parallel to further B-cell proliferation) by point mutation at rate *ρ*_*i*_, until the emergence of a mutated B-cell with antibodies at distance *i* − 1.

The clone size *n*_*i*_ = *n*(*m, i, R*) and mutation rate *ρ*_*i*_ = *ρ*(*m, i, R*) for DNA length *m* are *i*-and-*R*-dependent decision variables of the model. The best such pair maximizes the probability of survival of the host until an antibody at distance *i* −1 is created. Letting *S*_*opt*_(*m, i, R*) be this maximal survival probability, the global survival probability under optimal proliferation/mutation management is 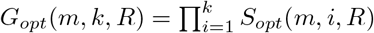. The alternative standard proliferation/mutation management, in which the same pair (*n, ρ*) is applied in each cycle, will be compared in Section 8 with optimal management.

More explicitly, standard proliferation/mutation management is defined (for given *m*) in terms of two “standard” parameters *n*_*sta*_(*m*) and *ρ*_*sta*_(*m*). At all distances *k* and against every risk rate *R*, B-cells are proliferated until a clone of size *n*_*sta*_(*m*) has been created, followed by point mutation at rate *ρ*_*sta*_(*m*). The particular choice to be adopted is the best pair (*n*_*sta*_(*m*) = *n*(*m*, 1, 0.0001), *ρ*_*sta*_(*m*) = *ρ*(*m*, 1, 0.0001)) to apply at distance *k* = 1 against risk *R* = 0.0001, for reasons to be clarified in Section 8. The survival probabilities *S*_*sta*_(*m, i, R*) and *G*_*sta*_(*m, k, R*) are defined accordingly, as functions of distance and risk.

### The clone size to onset mutation

As will be shown in Subsection 5.1, based on Agur et al., letting the clone size to onset mutation at distance *k* = 1 be *n* = *m* is essentially optimal. Furthermore, the best choice of basic strategy pair to apply at distance *i* is the best pair to apply at distance 1, had the DNA length *m* been 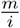 instead. Therefore, the extent of *m*-values to be analyzed should cover (at least) the range from 200*/*6 ≈33 to above 260.

The survival advantage of distance-dependent proliferation/mutation strategies has been established by Merkenschlager et al. (2025) in a study that combines modelling, experimental and simulation work. The current study focuses on mathematical modelling and optimization, confirming the optimality of distance-dependent proliferation/mutation management, while also revealing that management that ignores risk and distance can perform nearly as well as the optimal management.

### Organization

Section 2 describes the analysis by Agur et al. of the (superior but biologically infeasible) bang-bang management and the (*n, ρ*)-strategy, for recognition at distance *k* = 1. Section 3 evaluates the probability that the host will reach alive via proliferation the creation of a clone of size *n* of a B-cell. This section also introduces the notion of *survival coefficient* as a statistic that conveniently quantifies survival probability. Section 4 evaluates the probability that the host will reach alive via proliferation/mutation management the creation of a B-cell with higher affinity, starting from a clone of size *n* of the current B-cell. Special mathematical techniques are applied to solve a chaotic system of equations. Section 5 justifies (for *k* = 1) the assignment of the DNA length *m* as clone size *n* to onset mutation, and Section 6 extends the clone size assignment to general *k*. This section also evaluates the global survival probability under the optimal proliferation/mutation management, followed by the evaluation in Section 8 of the global survival probability under standard management, concluding that these survival probabilities are close. Subsection 7 evaluates the global survival probability under the bang-bang proliferation/mutation management, and concludes that the optimal and standard management are nearly as effective. Section 9 compares by simulation the performance of optimal management (based on the inhibitory working assumption that once a higher affinity antibody has been created the clone of B-cells that generated it ceases to be active) with its non-inhibitory counterpart. Section 10, Conclusion, summarizes the findings of the mathematical model of this study on the proliferation and mutation factors that govern pathogen recognition by the immune system. The Appendix documents the computations reported in an Excel data file attached as electronic supplementary material. Data for all tables in the text is taken from this Excel data file.

## 2 Recognition at distance *k* = 1 revisited. The bang-bang proliferation/mutation management

The setting for the current work is the analysis by Agur et al. (1991/2) of recognition at distance *k* = 1, which consisted of two steps, a purely theoretical step and a biologically feasible step. In the purely theoretical step the authors allowed the host to “know” at every moment the clone size *n* and to be able to set and change B-cell mutation rate at will. As proved to be optimal, purely proliferate B-cells until reaching a certain clone size *n*_0_ (slightly smaller than *m*) and then apply infinitesimally an infinite mutation rate to the *mn*_0_ locations to create one such mutation. If it is the correct mutation (with probability 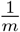), the process is finished and the host survived. If it is an adverse mutation, clone size decreases to *n*_0_ −1 and the mutation rate reverts to zero, waiting for a proliferation. In other words, geometrically many (on the average *m*) such “bang-bang” attempts are made to correct the single error. The advantage of a higher clone size *n*_0_ is that these attempts will be done more often, but it takes longer to create it.

This theoretical bang-bang proliferation/mutation management is not biologically feasible. Once started, hypermutation should keep a steady rate. If the rate is too high, clone size will experience a negative drift and the antibody repertoire may become empty, leading to death. In the biologically feasible step, the authors showed that for a mutation rate below but close to the drift-zero rate, applied steadily upon reaching some clone size, the survival probability is only slightly smaller than the upper bound provided by bang-bang management.

The current study extends this comparison from recognition at distance *k* = 1 to higher *k*, and shows that bang-bang proliferation/mutation management would have provided no real advantage. In fact, as will be shown in Subsection 7, over the entire parameter range covered in this study, bang-bang management reduces the probability of death by less than 1.1% with respect to optimal management (as defined in the Introduction). See the paragraph following Table 3.

### Rationale for the application of basic strategies (*n, ρ*)

Resorting to more general clone-size-dependent, progressively changing mutation rate could in principle reduce the probability of death, but not beyond bang-bang management. Since the latter does not provide real advantage, attention is restricted to the basic strategy (*n, ρ*) (applied in every affinity-improving cycle) that applies pure proliferation, and then a fixed mutation rate *ρ* once some threshold clone size *n* has been reached. Bang-bang management has a catalyst role in Agur et al. and in the current study. It provides an upper bound to performance against which to compare biologically feasible rules, but it is not intended as management proper.

## 3 Surviving proliferation

This section presents the probability *T* (*n*; *R*) that the host (via B-cell proliferation at rate *λ*, against a pathogen that kills the host with rate *r* = *λR*) reaches alive the creation of a clone of *n* B-cells, starting from one B-cell. *T* is clearly a decreasing function of *n*, starting near 1 if *R* is small enough,and tending to 0 at ∞.

### Markovian assumption

Clone size is modelled as a sub-stochastic continuous time Markov chain. This means that clone size will stay *n* until the minimum of two latent independent exponentially distributed random times, proliferation and death. The only possible transition from state *n*, to state *n* + 1, has the probability 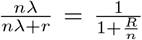 that proliferation occurs first. The complementary probability corresponds to death. To understand the term *nλ* in the transition probability from state *n* to state *n* + 1, proliferation time is the earliest of *n* independent exponentially distributed single-unit latent proliferation events.

The survival function *T* is thus

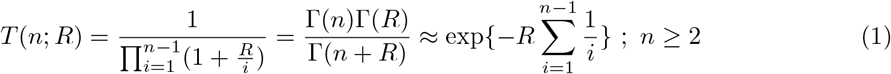

For numerical purposes, the recursive product form should be preferred, as the Gamma function grows too fast. However, an almost exact closed-form formula may be developed for the function *T*, as long as *R* is small enough. The key to the formula is Euler’s approximation to the harmonic sum

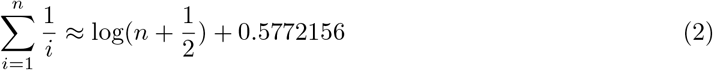

from which

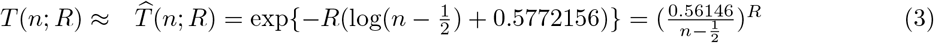

### 3.1 Probability of death and the survival coefficient

Motivated by the simplicity of 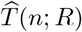, a generic survival probability *S* will be quantified in terms of the *survival coefficient* 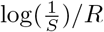. For the approximate survival probability 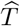 in the proliferation stage, this coefficient is 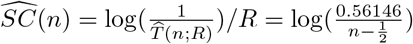, free of *R*.

Proliferation survival coefficients have values in [4, 6.3] with 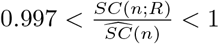 throughout the range of interest, 10^*−*4^ ≤*R* ≤10^*−*2^ and 30 ≤*m*≤ 300. *SC*(*n*; 0.0001) and 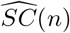 agree to three decimal places.

#### A small probability of death is nearly proportional to the survival coefficient

The probability of death 1−*S*, complementary to the survival probability *S*, is 1−*S* ≈ − log(*S*) = *SC* \**R*.

The survival coefficient admits an alternative *certainty equivalent* interpretation. Consider an event such as the creation of a clone of B-cells of some size or a favorable mutation, or global survival. With “unit” time consisting of the mean time to B-cell division, let *Z* be the exponentially distributed latent killing time by the pathogen (with mean 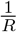) and let *τ* be the (latent) duration time until the event occurs, in the absence of the risk posed by the pathogen. Then the probability of surviving the event is *S* = *P* (*Z > τ*) = *E*[*P* (*Z > τ* |*τ*)] = *E*[exp{−*R* * *τ*}]. On the other hand, it is *S* = exp{−*R* * *SC*}, the inverse of 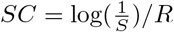. I.e.,

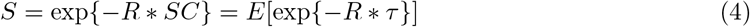

This equality identifies *SC* as the certainty equivalent^1^ of the latent random duration *τ* : *SC* is the deterministic duration of the immunological process that yields the same survival probability as the stochastic duration time *τ* (where both refer to duration in the absence of risk from the pathogen). This temporal interpretation of the survival coefficient will play a role in Section 7.

#### Additivity property

If the host undergoes independent risk events in tandem, the overall survival coefficient is the sum of the individual survival coefficients. This is specifically relevant for the analytical model adopted, where the creation of an antibody with higher affinity is followed by inhibition of the proliferation/mutation activity of lesser affinity B-cells, inducing independence of the successive risk events. Comparison with a non-inhibitory model will be performed by simulation in Section 9.

## 4 Surviving mutation until creating an antibody with higher affinity

This section introduces mutation rate *ρ* into the model. The section’s preamble and Subsections 4.1and 4.2 follow standard mathematical derivations similar to those of Section 3. Subsection 4.3, mathematically more elaborate, proposes an algebraic method to solve a chaotic system of equations. The time until the next mutation at clone size *n* is exponentially distributed with rate *mnρ*, as the minimum of the *mn* independent mutation times of the individual base pairs of the DNA sequence encoding the variable binding region, in each of the *n* antibodies. As usual in next-event Markov modelling, the latent proliferation, mutation and death times are independent, and the earliest “wins”.

Formally, let *A*(*n*; *m, k, R, λ, ρ*) be the probability that the host with DNA length *m*, via B-cell proliferation at rate *λ* and point mutation at rate *ρ*, against the pathogen that kills the host with rate *r*, reaches alive the creation of one B-cell that produces antibodies at distance *k*− 1 from the antigen, starting from a clone of *n* B-cells with antibodies at distance *k* from the antigen.

As will be seen in the sequel (Eq. 8), the function *A* depends on *k* and *m* only via their ratio, the *effective DNA length* 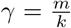 that generalizes the notion of DNA length *m* for *k >* 1, in a sense to be explained in Subsection 6. Mutation rate, as represented by the point mutation rate *ρ*, can (and will) be alternatively represented by the *relative mutation rate* 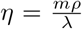, that sets the drift of the clone size process as long as the host is alive. For *η* = 1 the drift is zero, i.e., the proliferation rate *λ* equals the global rate of mutation *mρ*. For *η <* 1 clone size tends to increase.

The function *A* will henceforth be *A*(*n*; *γ, R, η*). This function is increasing, tending to 1 at ∞. *R* and *γ* are considered as given, while *η* is an optimization tool. Let *B*(*n*; *γ, R*) = max_*η*_ *A*(*n*; *γ, R, η*) keep *n* general, but optimize over *η* the survival probability in the mutation stage, starting from a clone of *n* B-cells.

The previous section evaluated the probability *T* (*n*; *R*) that the host reaches alive, via pure proliferation, the clone of *n* B-cells that are the starting point of the proliferation/mutation process analyzed in the current section. The product *W* (*n*; *γ, R*) = *T* (*n*; *R*)*B*(*n*; *γ, R*) is the maximal possible probability that the host reaches alive proliferation to clone size *n* and the creation by proliferation/mutation of one B-cell with antibodies at distance *k* −1 starting from one B-cell with antibodies at distance *k*. The optimal survival probability under the basic strategy at distance *k* is max_*n*_ *W* (*n*; *γ, R*).

### 4.1 Markovian analysis

Consider a clone of *n* proliferating and mutating B-cells with antibodies at distance *k* from the antigen. If the next change is a proliferation (the probability of this event is 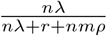), the clone size increases to *n* + 1. If the next change is a favorable mutation (the probability of this event is 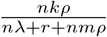), a B-cell with antibodies at distance *k* −1 has been created, and the host has survived this stage. The next change may be death (with probability 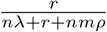). With the complementary probability 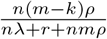, the clone size with antibodies at distance *k* decreases to *n* − 1.

#### Remark about adverse mutations

With the above mentioned complementary probability, an antibody-like protein at distance *k* + 1 has been created. It may be a lost protein or it may be a legitimate immunoglobulin recognized as having a lesser affinity with the antigen. The possible contribution of this antibody to survival will be ignored throughout. Hence, the survival probabilities to be analyzed and evaluated are lower bounds to those in “real life”. It is reasonable to postulate that the potential contribution of these adverse mutations is very minor.

A less negligible potential contribution stems from the clone of B-cells with antibodies at distance *k*, that can still continue to proliferate and mutate after a B-cell at distance *k* −1 has been created. Their overall contribution is to add to the growing clone or other immunoglobulin units at distance *k*− 1 a random number of B-cells, accelerating proliferation. The conservative practice of ignoring this effect circumvents the need for specifying what actually happens to these proteins in situ, continue proliferation and mutation or undergo apoptosis or regulation. As explained in Section 3, the analysis of strategies may ignore this issue. Simulations presented in Section 9 provide some further insight.

### 4.2 Recursive equation for the survival probability *A* in the mutation stage

Summarizing the Markovian analysis, and keeping parameters of *A* tacit,

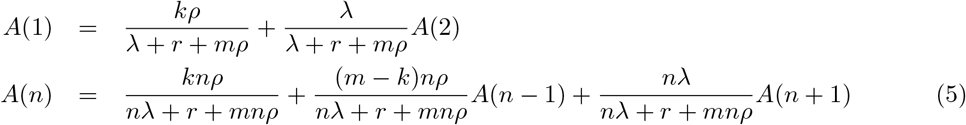

which can be expressed in terms of 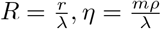 and 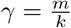 as

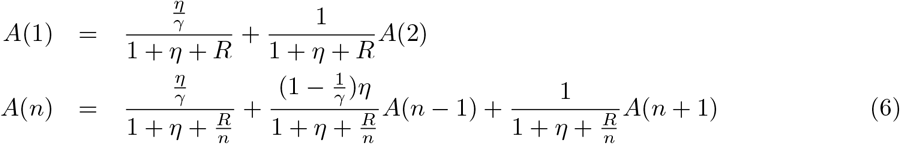

leading to the recursive formula

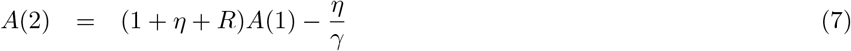

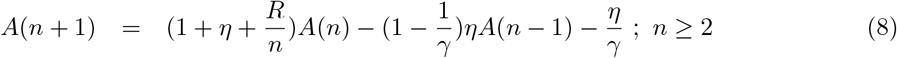

with boundary condition

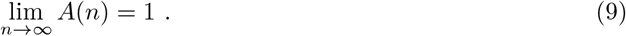

### 4.3 Evaluation of the survival probability *A* in the mutation stage

The evaluation of the function *A* is not as straightforward as that of the function *T*.

If *A*(1) is assigned an arbitrary positive value, recursion (7,8) yields a unique solution *A*. I.e.,

#### The family *A* of solutions of Eqs. (7,8)

There is a family *A* of solutions of Eqs. (7,8), parameterized by the initial value *A*(1). As will be shown, only one element *A*_0_ of *A* satisfies condition (9).

Clearly, there exist two sequences *α* and *β*, that do not depend upon *A*(1), such that the solution of Eqs. (7,8) for arbitrarily fixed *A*(1) is expressible as 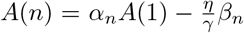. Their initial values are *α*_1_ = 1, *α*_2_ = 1 + *η* + *R, β*_1_ = 0, *β*_2_ = 1. Plugging this representation of *A* into (8) leads to

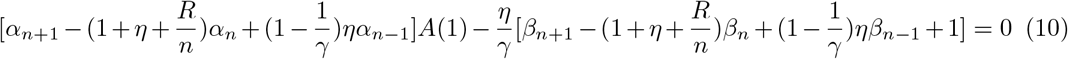

Since expression (10) is zero for arbitrary *A*(1), each of the two expressions in square brackets must vanish too. Hence, with the two first entries of *α* and *β* as determined above, *α* and *β* can be recursively built as

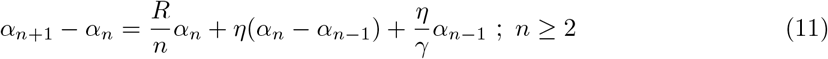

and

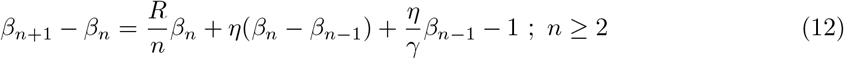

#### Properties of the sequence *α*

Recursion (11) provides an inductive proof that *α* is a positive increasing sequence, necessarily unbounded because its increment *α*_*n*+1_ − *α*_*n*_ exceeds the increasing sequence 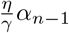.

For any *A*_1_, *A*_2_ ∈ A with *A*_2_(1) *> A*_1_(1), the difference *A*_2_(*n*) − *A*_1_(*n*) = *α*(*n*)(*A*_2_(1) − *A*_1_(1)) is positive, strictly increasing, diverging to ∞.

#### Bifurcation behavior

The survival probability *A*_0_ is increasing and satisfies (9), with a precise choice *A*_0_(1) of *A*(1). For *A*_1_(1) *< A*_0_(1), the function *A*_1_ (of the 1^*st*^ kind) reaches a maximal value below 1 and diverges to −∞, while for *A*_2_(1) *> A*_0_(1), the function *A*_2_ (of the 2^*nd*^ kind) crosses 1 and diverges to +∞.

Figure 1 (left) displays such a pair of functions, with initial values *A*_1_(1) and *A*_2_(1) differing by 10^*−*15^. The bifurcation behavior is apparent. Figure 1 (right) will be explained at the end of this section.

**Figure 1.**
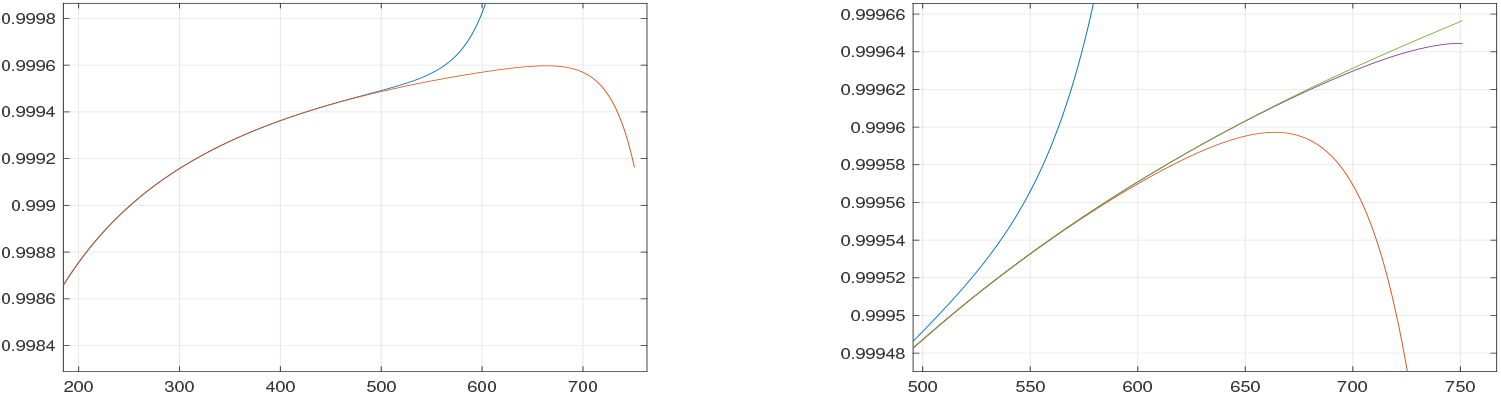
Left: two solutions (red - 1^*st*^ kind, blue - 2^*nd*^ kind) of the recursive equation (7,8), showing bifurcation behavior. DNA length *m* = 250, relative mutation rate *η* = 0.95, risk *R* = 0.001, distance *k* = 1. The two initial values *A*(1) differ by 10^*−*15^. Right: Zoom of the graph on the left with the addition of two functions *A*_1_ and *A*_2_ solving (13) with *N* = 750.

To motivate the solution for *A*_0_ introduced in the next paragraph, the function *A*_1_ to be defined is of the 1^*st*^ kind, the function *A*_2_ of the 2^*nd*^ kind.

#### Two systems of linear equations for *A*_0_

A numerically stable method to solve Eq. (7,8) respecting (9), is to view the pair as a system of linear equations for *A*_0_(*n*), 1≤*n*≤ *N* + 1, rather that as a recursion, where *N* amply exceeds *m* (e.g., *N* = 10*m*). Re-writing (8) as the linear equation

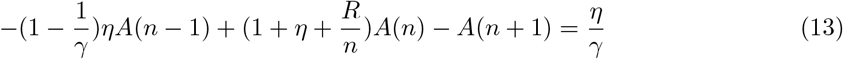

the linear system of equations is *QA*_0_ = *V*, where the tri-diagonal band matrix *Q*, of size *N* +1× *N* +1, has diagonal terms 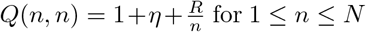 for 1 ≤ *n* ≤ *N*. The subdiagonal has equal entries 1 and the superdiagonal has equal entries 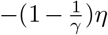. The column vector *V* has equal entries 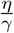 for 1 ≤ *n* ≤ *N*.

This specifies *N* equations for *N* + 1 variables. It remains to specify the last row of *Q* and last entry in *V*. This last entry is *V* (*N* + 1) = 0. As far as *Q*(*N* + 1, ·) is concerned, there are two versions, with solutions *A*_1_ and *A*_2_.

The first version, with *Q*(*N* +1, *N*) = −1 and *Q*(*N* +1, *N* +1) = 1, makes *A*_1_ have zero derivative near *N*, and as such is of the 1^*st*^ kind, a lower bound to *A*_0_.

The second version, with *Q*(*N* + 1, *N*) = −2 and *Q*(*N* + 1, *N* −1) = *Q*(*N* + 1, *N* + 1) = 1, makes *A*_2_ have zero second derivative near *N*, and as such is of the 2^*nd*^ kind, an upper bound to *A*_0_.

If *A*_1_(*m*) and *A*_2_(*m*) are close enough to each other, their common value provides *A*_0_(*m*).

The bifurcating pair of the recursion version is illustrated in Figure 1 left and as the two outer functions in Figure 1 right. The pair *A*_1_, *A*_2_ of the two systems of linear equations is displayed as the two inner functions in Figure 1 right, practically identical except for clone sizes close to *N* = 750 (out of the range of interest, as intended). At *n* = *N, A*_1_ is flat (zero derivative) and *A*_2_ is linear (zero second derivative, turning from concave to convex).

This approach via the two systems of linear equations is called for when the clone sizes of interest fall beyond the bifurcation of the recursive approach. As will be seen in Section 8, this is strongly the case for the analysis of standard proliferation/mutation management.

## 5 The basic strategy for *k* = 1 and the choice of the DNA length *m* as clone size to onset mutation

The proposed model requires to fix the clone size to onset the mutation stage. Emphasis will be placed on the choice of the DNA length *m* as candidate clone size at distance *k* = 1. The near optimality of this choice will be justified in Subsection 5.1. Implications for general *k* are handled in Section 6.

The entry of interest in the vector *A*_0_ to identify the basic strategy, is *A*_0_(*m*). As stated at the beginning of Subsection 4.2, the parameters of *A* have been kept tacit. *A*_0_(*m*) is really *A*_0_(*m*; *m, R, η*), considered now a function of *η*. This function is maximized over *η*, to obtain the best choice *η* = *η*(*m*, 1, *R*). In practice, *η* has been restricted to be a multiple of 0.001. The basic strategy at *k* = 1 is to proliferate the B-cells to clone size *m* and then apply (in parallel to further proliferation) mutation at relative mutation rate *η*(*m*, 1, *R*) until a favorable mutation emerges. The survival probability under this strategy is *W* (*m*; *m, R*) = *T* (*m, R*)*A*_0_(*m*; *m*, 1, *η*(*m*, 1, *R*)) represented in Table 1 by the survival coefficient *SC*_*prol*_ + *SC*_*mut*_. Table 1 focuses on *k* = 1 and displays the best choice of relative mutation rate *η*, and the corresponding proliferation/mutation survival coefficients *SC*_*prol*_ and *SC*_*mut*_ for a menu of values of *m*.

**Table 1:** For clone size *n* = *m*, best choice of relative mutation rate *η* (taken as a multiple of 0.001) for *k* = 1, survival coefficient 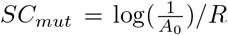 in the mutation stage and survival coefficient 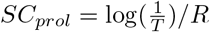 in the proliferation stage. *SC*_*tot*_ is the sum of the two. Entries in bold type lead to the entries in bold type in Table 2.

| *R* | 0.01 | *SC* | *SC* | *SC* | 0.001 | *SC* | *SC* | *SC* | 0.0001 | *SC* | *SC* | *SC* |
|---|---|---|---|---|---|---|---|---|---|---|---|---|
| *m* | *η* | *mut* | *prol* | *tot* | *η* | *mut* | *prol* | *tot* | *η* | *mut* | *prol* | *tot* |
| 30 | 0.830 | 1.045 | 3.954 | 4.999 | 0.760 | 1.069 | 3.961 | 5.030 | 0.697 | 1.094 | 3.962 | 5.055 |
| 48 | 0.900 | 1.022 | 4.430 | 5.452 | 0.851 | 1.034 | 4.437 | 5.471 | 0.805 | **1.046** | **4.438** | **5.484** |
| 60 | 0.925 | 1.015 | 4.655 | 5.670 | 0.885 | 1.025 | 4.662 | 5.687 | 0.846 | **1.032** | **4.663** | **5.696** |
| 80 | 0.950 | 1.009 | 4.945 | 5.954 | 0.920 | 1.016 | 4.952 | 5.968 | 0.889 | **1.021** | **4.953** | **5.974** |
| 120 | 0.973 | 1.004 | 5.352 | 6.356 | 0.957 | 1.009 | 5.360 | 6.369 | 0.936 | **1.011** | **5.360** | **6.372** |
| 200 | 0.989 | 1.000 | 5.865 | 6.865 | 0.982 | 1.005 | 5.872 | 6.877 | 0.973 | 1.005 | 5.873 | 6.878 |
| 210 | 0.990 | 1.000 | 5.914 | 6.914 | 0.983 | 1.004 | 5.921 | 6.925 | 0.975 | 1.005 | 5.922 | 6.927 |
| 220 | 0.990 | 1.000 | 5.960 | 6.960 | 0.984 | 1.004 | 5.968 | 6.972 | 0.977 | 1.005 | 5.969 | 6.973 |
| 230 | 0.991 | 0.999 | 6.005 | 7.004 | 0.985 | 1.004 | 6.012 | 7.016 | 0.979 | 1.005 | 6.013 | 7.018 |
| 240 | 0.992 | 0.999 | 6.048 | 7.047 | 0.986 | 1.004 | 6.055 | 7.059 | 0.981 | **1.004** | **6.056** | **7.060** |
| 250 | 0.992 | 0.999 | 6.089 | 7.088 | 0.987 | 1.004 | 6.096 | 7.100 | 0.982 | 1.004 | 6.097 | 7.101 |
| 260 | 0.993 | 0.999 | 6.128 | 7.127 | 0.988 | 1.003 | 6.135 | 7.138 | 0.984 | 1.004 | 6.136 | 7.140 |
| 300 | 0.995 | 0.998 | 6.271 | 7.269 | 0.990 | 1.003 | 6.279 | 7.282 | 0.988 | 1.003 | 6.279 | 7.283 |

### Proliferation is riskier than mutation

Table 1 illustrates that the probability of death during the mutation stage decreases in *m* and is approximately *R*, while the probability of death during the proliferation stage increases in *m* and is between 4 and 6.3 times *R* (in the displayed range). The probability of death during the basic proliferation/mutation cycle increases in the DNA length *m*.

### 5.1 The nearly optimal choice of clone size *n* = *m* for *k* = 1

Switching now to general clone size *n*, but keeping the interpretation of *A*_0_ as a function of *η*, this function is maximized to obtain *B*(*n*; *m, R*) = max_*η*_ *A*_0_(*n*; *m, R, η*). The objective function (in terms of *n*) becomes *W* (*n*; *m, R*) = *T* (*n*; *R*)*B*(*n*; *m, R*), as introduced at the beginning of Section 4. The current subsection deals with the choice of the clone size *n*.

The factor *T* (of *W*) decreases in *n* (survival to clone size *n* + 1 requires survival to clone size *n*, see (1)) while the factor *B* increases with *n* (at higher clone size, proliferation occurs more often). Their product *W*, to be maximized, need not be unimodal in principle.

The survival probability values *W* (*n*; *m, R*) are transformed as suggested in Subsection 3.1 into their corresponding survival coefficients 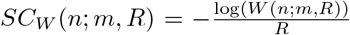. The first task will be to show analytically (in (14)) that search for the minimum of *SC*_*W*_ may be restricted to clone sizes *n <* 3*m*. The second task, performed numerically and displayed in Figure 2, is to show the near optimality of the choice of the DNA length *m* as clone size to onset mutation.

**Figure 2.**
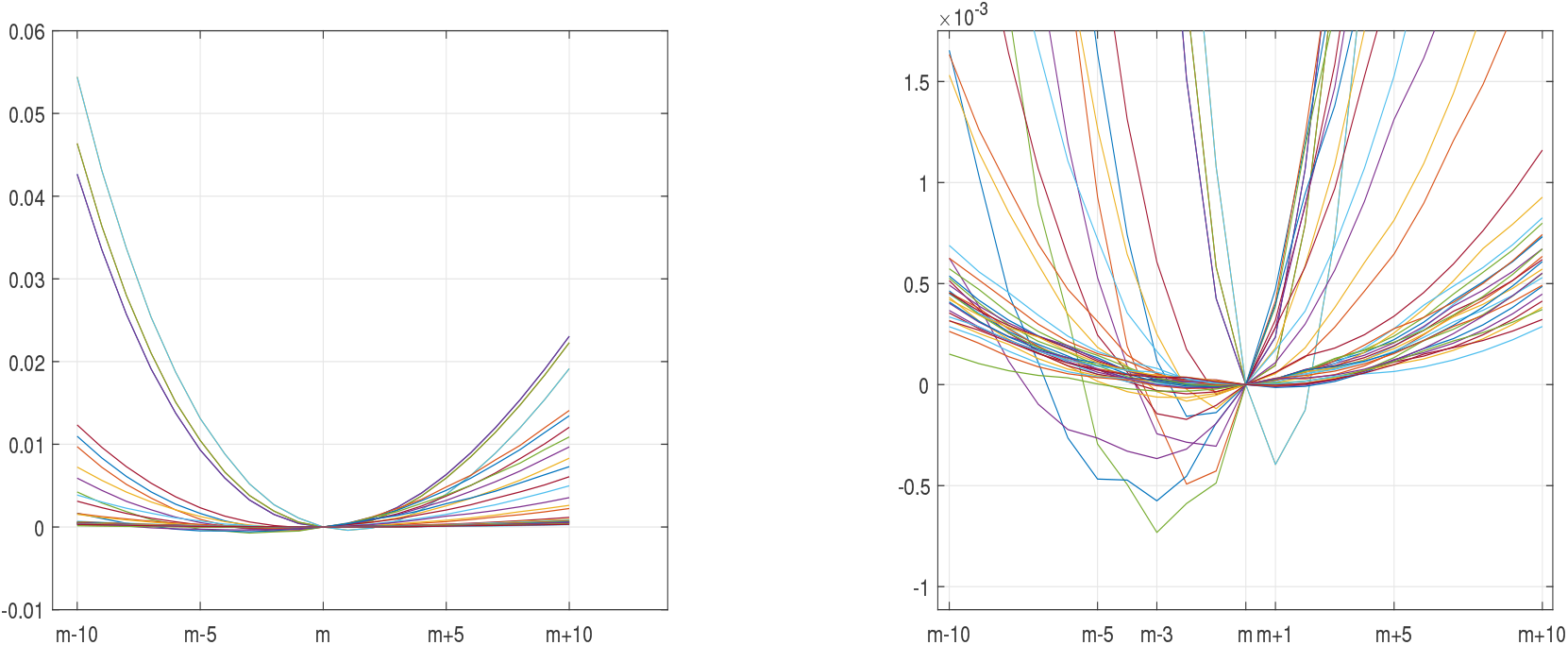
: The choice of clone size *n* for *k* = 1. The figure on the left displays (for the choices of *R* and *m* in Table 1 plotted together, and clone size *n* between *m* −10 and *m* + 10) the excess function *D*(*n*; *m, R*) of the basic survival coefficient over its value at clone size *n* = *m*. The figure on the right, a vertical zoom of the figure on the left, indicates that the optimal clone size *n* is between *m*− 3 and *m* + 1. It also indicates that the minimal survival coefficient differs from the survival coefficient at clone size *m* by less than 10^*−*3^.

Identifying *T* as 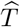 (see (3)), *SC*_*prol*_(*n*) = log(*n* * *const*) and *SC*_*prol*_(3*m*) = *SC*_*prol*_(*m*) + log(3).

As for *SC*_*mut*_(*m*), it can be seen in Table 1 that it is at most 1.0936 *<* log(3). Thus,

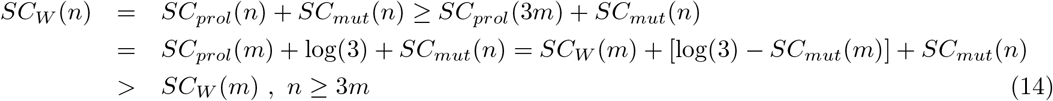

Since *SC*_*W*_ (*n*) *> SC*_*W*_ (*m*) for all *n* ≥3*m*, the search for a minimal *SC*_*W*_ (*n*) can be restricted to *n <* 3*m*.

Let *D*(*n*; *m, R*) = *SC*_*W*_ (*n*; *m, R*) −*SC*_*W*_ (*m*; *m, R*) be the excess of *SC*_*W*_ (·; *m, R*) over the value at *n* = *m*. Figure 2 (left) displays together all 39 functions *D* (three values of *R* times thirteen of *m* as in Table 1), for clone size *n* between *m* −10 and *m* + 10, after having checked numerically that these functions *D* are indeed unimodal in the wider range of *n* values up to 3*m*.

Tables 1 and 2 set the value of *SC*_*W*_ (*m*; *m, R*) as exceeding 5 over the entire range. Figure 2 (right) illustrates that the optimal choice of clone size *n* can reduce the survival coefficient by less than 0.001 with respect to the choice *n* = *m*. Therefore,

**Table 2:** Global survival coefficients under optimal proliferation/mutation management for *m*-derived clone sizes and risk *R* = 0.0001. Entries in bold type are derived from the entries in bold type in Table 1.

| *m* \\*k* | 1 | 2 | 3 | 4 | 5 | 6 |
|---|---|---|---|---|---|---|
| 60 | 5.696 | 10.751 | 15.468 | 19.975 | 24.349 | 28.640 |
| 120 | 6.372 | 12.067 | 17.381 | 22.437 | 27.301 | 32.018 |
| 200 | 6.878 | 13.070 | 18.867 | 24.389 | 29.704 | 34.852 |
| 210 | 6.927 | 13.167 | 19.011 | 24.579 | 29.938 | 35.131 |
| 220 | 6.973 | 13.259 | 19.148 | 24.761 | 30.163 | 35.398 |
| 230 | 7.018 | 13.347 | 19.279 | 24.934 | 30.378 | 35.653 |
| **240** | **7.060** | **13.432** | **19.405** | **25.101** | **30.585** | 35.899 |
| 250 | 7.101 | 13.513 | 19.526 | 25.261 | 30.783 | 36.135 |
| 260 | 7.140 | 13.591 | 19.642 | 25.415 | 30.974 | 36.362 |
| 300 | 7.283 | 13.875 | 20.067 | 25.978 | 31.673 | 37.196 |

**Table 3:** *R*-free approximate global survival coefficients under bang-bang proliferation/mutation management.

| *m*\\*k* | 1 | 2 | 3 | 4 | 5 | 6 |
|---|---|---|---|---|---|---|
| 60 | 5.638 | 10.549 | 15.021 | 19.171 | 23.062 | 26.737 |
| 120 | 6.348 | 11.986 | 17.202 | 22.113 | 26.784 | 31.256 |
| 200 | 6.865 | 13.028 | 18.775 | 24.224 | 29.439 | 34.463 |
| 210 | 6.915 | 13.127 | 18.924 | 24.424 | 29.691 | 34.766 |
| 220 | 6.962 | 13.221 | 19.066 | 24.614 | 29.930 | 35.054 |
| 230 | 7.007 | 13.311 | 19.202 | 24.796 | 30.158 | 35.329 |
| 240 | 7.050 | 13.398 | 19.332 | 24.970 | 30.376 | 35.592 |
| 250 | 7.091 | 13.480 | 19.456 | 25.136 | 30.585 | 35.844 |
| 260 | 7.130 | 13.560 | 19.576 | 25.296 | 30.786 | 36.086 |
| 300 | 7.274 | 13.849 | 20.011 | 25.879 | 31.517 | 36.966 |

#### Paradigmatic choice of clone size to onset mutation

For *k* = 1, the clone size to onset mutation is henceforth identified as the DNA length *m*. For simplicity, with some abuse of language, the corresponding nearly optimal strategies and parameters will be termed *optimal*.

## 6 The choice of clone size 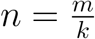 for general *k*

Consistent with recursion (7,8), where *m* and *k* appear only through their ratio 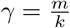, if the clone size to onset mutation at distance *k* = 1 is fixed as *m*, the appropriate clone size to onset mutation at distance *k* is an integer next to *γ*. As was seen in Subsection 5.1, clone size can be perturbed by 1 or 2, so that with the properly adjusted mutation rate, survival probability remains essentially the same. It is accurate enough to approximate the survival coefficient of the basic strategy as the weighted linear interpolation between the two survival coefficients *SC*(⌊*γ* ⌋, *R*) and *SC*(⌈*γ*⌉, *R*). A nearly optimal *η*(*m, k, R*) is obtained from *η*(·, 1, *R*) accordingly.

Table 2 displays for risk *R* = 0.0001 the global survival coefficients. The entries for *k* = 1 are taken from the last column of Table 1. Based on 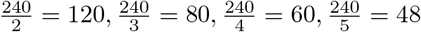, the entries for *m* = 240 are cumulative sums of the last column of Table 1:

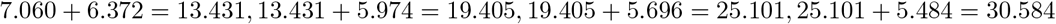

Under the working assumption of inactivity of lesser affinity B-cells, the basic strategy is applied and analyzed for each distance separately. As announced in the first paragraph of this section, and evident in the recursive equation (7,8), the function *A* depends on *m* and *k* only via their ratio, the effective DNA length *γ*. Hence, evaluation of *A* for distance *k* and DNA length *m* is equivalent to evaluating *A* for distance 1 and DNA length 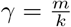. Thus, in order to evaluate the functions *A* for DNA lengths ranging from 200 to 260 and distance ranging from 1 to 6, it is enough to consider *k* = 1 but values of *m* ranging from 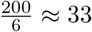 to 260.

### Remark on the working range of *R* values

It is worthwhile pointing out that for recognition at distance *k* = 6 and DNA length *m*≥ 200 against risk *R* = 0.01, the pathogen is very lethal, killing the host with probability over 36% (survival coefficient 36.362 in Table 2).

This is why the operative upper limit of *R* was not chosen higher.

## 7 Bang-bang proliferation/mutation management

The evaluation of survival coefficients under bang-bang management follows similar lines but is much simpler than the analysis above.

Agur et al. proved that the maximal survival probability at *k* = 1 is achieved by bang-bang proliferation/mutation management that alternates between immediate mutation of one amino-acid and waiting for a proliferation, until the mutation is favorable. This is not considered biologically feasible, but can provide an upper bound to the survival probability. The purpose of this section is to show, for general *k*, that the infeasible but hyper-optimal bang-bang management would have been only slightly more efficient than keeping steady mutation rates. As a result, there is no need to consider anything but steady mutation rates applied upon reaching some clone-size thresholds via pure B-cell proliferation. Agur et al. proved analytically for bang-bang management that the optimal threshold is ⌊ *m*(1 − *R*) ⌋. This is consistent with Figure 2 (right) and supportive of the paradigmatic choice of the DNA length *m* as the operative threshold.

Under the application of the bang-bang rule, with proliferation to clone size 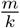 for distance *k*, the survival probability *S*_*bb,mut*_ in the mutation stage satisfies 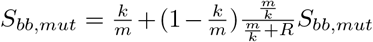 either produce a correct mutation (with probability 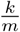) or else a wrong one, followed by one proliferation (with probability 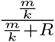)and repeat all over. Hence, 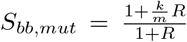. The bang-bang survival coefficient in the mutation stage

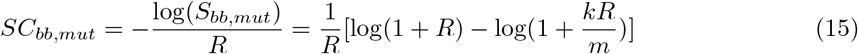

differs from 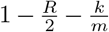 by less than 10^*−*4^ throughout the range of *m, k* and *R* adopted in this study.

Table 3 is based on the *R*-free approximation 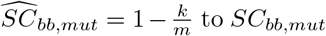. Thus, the exact figures may be smaller for *R* = 0.01 by 0.005 with respect to the reported values.

Table 3 displays the *R*-free approximate global survival coefficients of bang-bang management, cumulative sums of the basic survival coefficient approximations 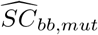 and 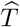,

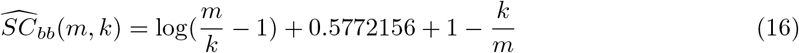

It can be observed that entries in Table 3 are only slightly smaller than those reported in Table 2 for *R* = 0.0001, that *exceed* those for bigger *R*. The bang-bang formulas provide adequate approximations to biologically feasible management.

The cumulative sums of the basic survival coefficients (16) reported in Table 3 provide adequately accurate formulas for the **optimal global survival coefficients**. No such simple representation has been found for the best choices of relative mutation rates *η* (see Table 1), that seem to require the heavy machinery to solve Eqs. (7,8,9).

Eq. (4) provides a temporal certainty equivalent interpretation of the survival coefficient. E.g., the entry 30.376 for DNA length *m* = 240 and affinity distance at recognition *k* = 5 in Table 3 establishes that if the immunological process had taken (in the absence of risk) *deterministic* duration time that is 30.376 times the mean time 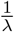 for proliferation, the probability of survival would have been the same as that induced by bang-bang or optimal management. At the commonly reported mean time for proliferation between 8 to 10 hours, this certainty equivalent duration time is somewhere between 10 to 12 days.

### Remark on the expected duration time of the immunological process

It may be of interest to append to this conceptual but somewhat unusual notion of typical time a more common one, the expected duration time of the immunological process. However, for optimal and standard management there is a positive probability that the immunological process will end with the depletion of the bank of B-cells with antibodies that recognize some affinity with the pathogen. This implies that the expected duration of the immunological process in the absence of risk is infinite. Thus, expected duration is not a meaningful objective, in contrast to the certainty equivalent.

Simple formulas can be derived for the expected duration time of bang-bang management, that comes out bigger than but very close to the survival coefficient. Details are omitted, for lack of relevance to biologically feasible management.

## 8 The standard proliferation/mutation management

The previous sections have optimized clone size and mutation rate, adapted to DNA length *m*, distance *k* and risk *R*. The current task is to show that there is no evolutionary need to adapt clone size and mutation rate to the distance *k* and certainly not to the generally unknown threat *R* posed by the pathogen. With this in mind, standard management, as outlined in the Introduction, fixes the clone size as *m* for all 1 ≤ *k* ≤ 6 and the relative mutation rate *η*, for each such *k* and *R* ∈ {0.01, 0.001, 0.0001} as the optimal choice *η*(*m*, 1, 0.0001) for *k* = 1 and *R* = 0.0001. The survival coefficient of proliferation is the same for all *k*, 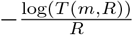. The survival probability in the proliferation/mutation cycle at distance *k* is readily identified as *A*_0_(*m*,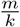, *R, η*(*m*, 1, 0.0001)).

### The choice of relative mutation rate as dictated by *R* = 0.0001

The optimal values displayed in Table 1 set the relative mutation rates *η* in terms of the DNA length *m*, which is legitimate, and of the risk *R*, which may not be. As displayed by Table 4, the high *η*-values designed for risk *R* = 0.01 yield high probabilities of death if applied against risk *R* = 0.0001, but the low *η*-values designed for risk *R* = 0.0001 yield probabilities of death only slightly higher than optimal if applied against risk *R* = 0.01 or 0.001. This pattern, quite pronounced for small DNA lengths, persists for bigger DNA lengths.

**Table 4:**
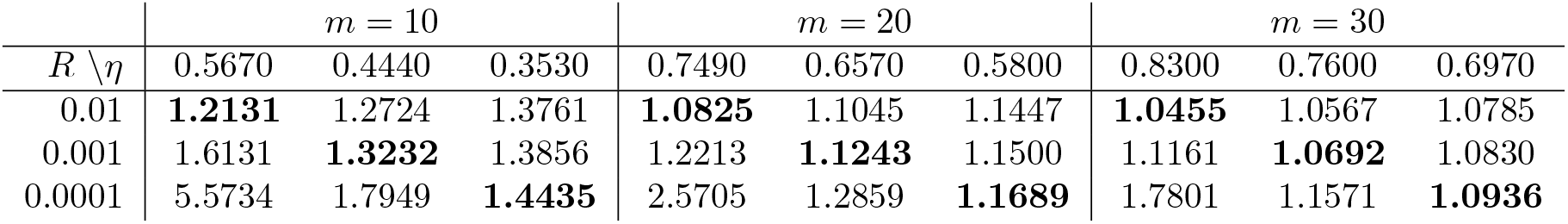
Survival coefficients for DNA lengths *m* = 10, 20, 30. Those corresponding to the optimal choices of relative mutation rate *η* appear in bold type. The optimal choice of *η* for *R* = 0.0001 performs reasonably well for bigger *R*, while the optimal choice for *R* = 0.01 yields high survival coefficients for smaller *R*.

The *η* values and optimal mutation survival coefficients (bold type) in Table 4 for *m* = 30 coincide with those reported in Table 1. Although not biologically meaningful, *η* values and survival coefficients are reported for *m* = 10 and 20 too, to highlight the more extreme differences experienced within the same pattern for small values of *m*. Each survival coefficient in regular type (*η* value that is not the best for the particular *R*) was computed explicitly for this table.

#### Remark about the evaluation of *A*_0_ by the two systems of linear equations

Evaluation of the function *A* in Eq. (8) involves clone size values *n* in the range from 1 to *γ*, and this range is generally in the initial stable regime of the chaotic recursion. Standard management applies clone size *m* for each 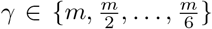. Hence, Eq. (8) needs to be applied for *n* in the range from 1 to 6*γ*, well past the bifurcation, as illustrated in Figure 1. This is where the method of linear equations, with large horizon *N*, shows its merits.

Table 5 displays for risk *R* = 0.0001 the global survival coefficients under standard management, for the menu of *m* values above. Optimal (Table 2) and standard (Table 5) survival coefficients are indeed close, at least for *R* = 0.0001. The next Section, comparing optimal, standard and bang-bang performance, will extend the comparison.

**Table 5:** Global survival coefficients under standard proliferation/mutation management for risk *R* = 0.0001.

| *m*\\*k* | 1 | 2 | 3 | 4 | 5 | 6 |
|---|---|---|---|---|---|---|
| 60 | 5.696 | 10.904 | 15.937 | 20.882 | 25.771 | 30.623 |
| 120 | 6.372 | 12.249 | 17.957 | 23.580 | 29.150 | 34.686 |
| 200 | 6.878 | 13.258 | 19.470 | 25.598 | 31.674 | 37.717 |
| 210 | 6.927 | 13.355 | 19.616 | 25.792 | 31.917 | 38.008 |
| 220 | 6.973 | 13.448 | 19.754 | 25.976 | 32.148 | 38.286 |
| 230 | 7.018 | 13.536 | 19.887 | 26.153 | 32.369 | 38.551 |
| 240 | 7.060 | 13.621 | 20.013 | 26.322 | 32.580 | 38.805 |
| 250 | 7.101 | 13.702 | 20.135 | 26.485 | 32.784 | 39.049 |
| 260 | 7.140 | 13.780 | 20.252 | 26.641 | 32.979 | 39.283 |
| 300 | 7.283 | 14.065 | 20.680 | 27.211 | 33.692 | 40.139 |

The standard proliferation/mutation management may be more evolution-efficient than meets the eye. The adopted mathematical model postulates that favorable mutations reduce distance one at a time. But suppose, to the contrary, that some mutation of a supposedly distance-6 B-cell could be so effective that it will produce a distance-2 B-cell directly. Standard management may be a safer response than applying initial clone size 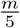.

As illustrated in Table 1, the global survival coefficients are decreasing functions of *R*, exceeding the bang-bang survival coefficients. To realize that the two types are close, it is enough to compare the bang-bang global survival coefficients with those for *R* = 0.0001. Table 2 is the biologically feasible counterpart of Table 3. For *m* ≥ 200, all corresponding entries in Tables 2 have ratios between 1 and 1.0113. As for a comparison between optimal and standard, the corresponding entries in Tables 2 and 5 have ratios between 1 and 1.0835.

To summarize the findings of this section,

**Keeping a steady relative mutation rate** *η* until an antibody of higher affinity has been created, is adequate enough. Letting *η* and the threshold clone size to onset mutation be fixed, independent of affinity, is almost as adequate. Hence, the immune system need not rely on the initial or progressive affinity of the antibodies towards the invading pathogen.

## 9 Simulation of the optimal and standard strategies with and without inhibition of lesser affinity B-cells

The goal of this section is to study the role of the working assumption that once a higher affinity antibody has been created, the clone of B-cells that generated it ceases to be active, by simulating proliferation and mutation in the absence of this assumption.

The optimal (Table 6) and the standard (Table 7) strategies are simulated for *m* = 60, 120 and 240, *k* = 3 and *R* = 0.01, 0.001. The relative mutation rates *η*, as in Table 1, are copied from the Excel file in the electronic supplementary material, columns C and D. Each of the 24 simulations was performed with sample size 10^6^.

**Table 6:** Simulation of the optimal proliferation/mutation management, applied at distance *k* = 3. The sample size is 10^6^ for each run. Model 1 represents management with proliferation and mutation inhibition and under Model 2, mutation and proliferation are active to the end. Average clone size reached at distance *j* is reported as *CS*_*j*_. The total number of mutations (adverse except for a few) is Mutat. The theoretical entries for optimal management are copied from the Excel file in the electronic supplementary material, columns Y and AF. For *m* = 240, Model 2 magnifies 5-fold the adverse mutations generated by Model 1. The probability of death in Model 1 is 11% higher than in Model 2.

| *m* | *R* | Model | $\log(\frac{1}{G})/R$<br>Optimal<br>empirical | Mutat | *CS*<sub>1</sub> | *CS*<sub>2</sub> | *CS*<sub>3</sub> | $\log(\frac{1}{G})/R$<br>Optimal<br>theoretical |
|---|---|---|---|---|---|---|---|---|
| 60 | 0.01 | 1 | 15.19 | 97 | 65 | 31 | 22 | 15.29 |
| 60 | 0.01 | 2 | 13.11 | 385 | 67 | 35 | 24 |  |
| 60 | 0.001 | 1 | 15.41 | 108 | 75 | 34 | 23 | 15.39 |
| 60 | 0.001 | 2 | 13.11 | 425 | 76 | 38 | 26 |  |
| 120 | 0.01 | 1 | 17.19 | 192 | 119 | 59 | 41 | 17.30 |
| 120 | 0.01 | 2 | 15.25 | 841 | 121 | 62 | 44 |  |
| 120 | 0.001 | 1 | 17.14 | 216 | 137 | 64 | 43 | 17.35 |
| 120 | 0.001 | 2 | 15.20 | 934 | 138 | 68 | 45 |  |
| 240 | 0.01 | 1 | 19.32 | 379 | 223 | 113 | 80 | 19.36 |
| 240 | 0.01 | 2 | 17.35 | 1826 | 228 | 117 | 82 |  |
| 240 | 0.001 | 1 | 19.25 | 433 | 261 | 124 | 83 | 19.40 |
| 240 | 0.001 | 2 | 17.46 | 2053 | 262 | 127 | 86 |  |

**Table 7:** Simulation of standard management, applied at distance *k* = 3. The sample size is 10^6^ for each run. Model 1 represents standard management with proliferation and mutation inhibition, and under Model 2, mutation and proliferation are active to the end. Average clone size reached at distance *j* is reported as *CS*_*j*_. The total number of mutations (adverse except for a few) is Mutat. The theoretical entries for standard management are copied from the Excel file in the electronic supplementary material, columns CJ and CQ. For *m* = 240, Model 2 magnifies 11-fold the adverse mutations generated by Model 1. The probability of death in Model 1 is between 20% and 23% higher than in Model 2.

| *m* | *R* | Model | $\log(\frac{1}{G})/R$<br>Standard<br>empirical | Mutat | *CS*<sub>1</sub> | *CS*<sub>2</sub> | *CS*<sub>3</sub> | $\log(\frac{1}{G})/R$<br>Standard<br>theoretical |
|---|---|---|---|---|---|---|---|---|
| 60 | 0.01 | 1 | 15.88 | 96 | 62 | 62 | 64 | 15.90 |
| 60 | 0.01 | 2 | 12.44 | 878 | 65 | 92 | 96 |  |
| 60 | 0.001 | 1 | 15.96 | 108 | 75 | 70 | 68 | 15.93 |
| 60 | 0.001 | 2 | 12.33 | 1183 | 78 | 147 | 171 |  |
| 120 | 0.01 | 1 | 17.90 | 191 | 116 | 119 | 124 | 17.93 |
| 120 | 0.01 | 2 | 14.60 | 1929 | 121 | 161 | 172 |  |
| 120 | 0.001 | 1 | 17.96 | 216 | 137 | 132 | 130 | 17.95 |
| 120 | 0.001 | 2 | 14.65 | 2326 | 139 | 202 | 223 |  |
| 240 | 0.01 | 1 | 20.02 | 376 | 220 | 229 | 241 | 19.90 |
| 240 | 0.01 | 2 | 16.65 | 4182 | 228 | 288 | 314 |  |
| 240 | 0.001 | 1 | 20.29 | 433 | 261 | 254 | 253 | 20.01 |
| 240 | 0.001 | 2 | 16.48 | 4815 | 263 | 330 | 354 |  |

These simulations were repeated for a model (Model 2) without inhibition of lesser affinity B-cells. All clones continue to proliferate and mutate, contributing occasionally higher affinity B-cells, that join the emerging clone at the corresponding distance from the pathogen’s antigens. All B-cells at a given such distance are considered as one growing “clone”, even if the mutated genes differ.

The death probabilities for the non-inhibitory model are, as expected, lower than under the inhibitory counterparts. However, this decrease (by 10% in the optimal setup and 18% in the standard setup) is accompanied by a sharp magnification of the trail of adverse mutations (5-fold and 11-fold respectively). Death probabilities show a mild 12% decrease from the former to the latter model - the two are comparable enough.

### Rationale for proliferation/mutation inhibition

These tables exhibit clearly the sharp magnification (5-fold for optimal management, 10-fold for standard management) of the trail of B-cells, predominantly adverse, left by the non-inhibitory versus the inhibitory model, to achieve a 10% to 28% reduction in probability of death. The search for apoptosis mechanisms (as in Opferman 2008 and Wensveen et al. 2012) or regulatory T-cells (such as the contribution of Yu Shan et al. (2023) to the field, which earned Brunkow, Ramsdell, and Sakaguchi the 2025 Nobel Prize in Medicine and Physiology) to dampen proliferation and mutation following the emergence of favorable mutations, could help ellucidate to some extent a process of evolutionary compromise.

## 10 Conclusion

The simplified, parsimonious mathematical model of this study avoids the specification of complex biological mechanisms. It attempts to explain with the partial support of the simulations above, with and without the application of affinity-driven strategies, with and without regulatory growth inhibition, that these factors do not affect survival probabilities in a critical way. Survival probabilities *G* are conveniently quantified by the survival coefficient *SC* = log(1*/G*)*/R*, nearly independent of *R* in the wide studied range, in which an unrecognized pathogen will kill the host after 37 days to 10 years on the average.

Survival probabilities and immunological response strongly depend on the (observed) length of the DNA sequence encoding the variable region of the antibody that binds to the antigen. The (unobserved) risk rate *R* posed by the pathogen and the initial affinity of the host’s B-cell antibodies to its antigen strongly affect survival probabilities, but propitiously have only a minor effect on determining the host’s response: the immunological response need not rely on the (unobserved) risk *R* nor on the initial and progressive affinity of the host’s B-cell antibodies to the pathogen throughout the maturation process. This allows the introduction of standard management, with survival probability close to the optimal survival probability, that applies a fixed mutation rate and onsets mutation at a fixed clone size, independent of affinity and risk.

## 11 Appendix The data depository, Excel file as electronic supplementary material

The tables through the paper report mutation rates and survival coefficients for a restricted menu of DNA length *m* values. The Excel file is a data depository that covers 60 ≤ *m* ≤ 300, 1 ≤ *k* ≤ 6 and the three choices 0.01, 0.001, 0.0001 of *R* in full, and 1 ≤ *m* ≤ 59 with *k* = 1 and the three choices 0.01, 0.001, 0.0001 of *R*, partially. Columns up to BK report numerical values evaluated via a Matlab program, and columns from BM onwards are Excel-based simple arithmetic operations.

Column B lists *m*. For distance *k* = 1 and the three choices of *R*, Columns C to E report best mutation rates, columns G to I survival coefficients in the mutation stage, columns K to M survival coefficients in the proliferation stage, columns P to R survival coefficients under bang-bang management and columns S to U under optimal management. Column N displays the harmonic sum, practically identical to column M.

Columns W to AB display for *R* = 0.01 and 1 ≤*k*≤ 6 the global survival coefficients under optimal management, columns AD to AI display the *R* = 0.001 counterparts, with columns AK to AP displaying the version with *R* = 0.0001.

Columns AR to AW, AY to BD and BF to BK display the corresponding survival coefficients in the mutation stage for standard management.

Columns BM to CF add previous columns to present the basic survival coefficients of standard management, and columns CH to DA add over these to present the global survival coefficients of standard management.

Columns DC to DO report the *R*-free approximations for the basic and global bang-bang survival coefficients.

## Acknowledgement

We are grateful to Zvia Agur, Gadi Mazor, Nechama Smorodinsky and Yariv Wine for comments and information.

## Footnotes

1 The term certainty equivalent is borrowed from Economics and Finance, where the certainty equivalent of a risky asset is a safe asset with the same expected utility.

## Notes

### Competing Interest Statement

The authors have declared no competing interest.

## References

[1] Agur, Z., Mazor, G. & Meilijson, I. Maturation of the humoral immune response as an optimization problem. Proceedings of the Royal Society of London, Series B: Biological Sciences 245 (1313), 147–150 (1991).

[2] Agur, Z., Mazor, G., & Meilijson, I. Mimicking the strategy of the immune system: insight gained from mathematics. Theoretical and Experimental Insights into Immunology, ed. A. Perlson, Springer Verlag Series on Cell Biology, 323–332. Springer-Verlag Berlin Heidelberg 1992.

[3] Merkenschlager, J., Pyo, A.G.T., Silva Santos, G.S.,Schaefer-Babajew, D., Cipolla, M., Hartweger, H., Gitlin, A.D., Wingreen, N.S. & Nussenzweig, M.C. Regulated somatic hypermutation enhances antibody affinity maturation. Nature 641, 495–420 (2025).

[4] Opferman, J.L. Apoptosis in the development of the immune system. Cell Death & Differentiation, 15, 234–242 (2008).

[5] Shan, Y., Zhao, J., Zheng, Y., Guo, Sh., Schrodi, S.J. & He, D. Understanding the function of the GABAergic system and its potential role in rheumatoid arthritis. Frontiers in Immunology 14, 2023. doi: 10.3389/fimmu.2023.1114350

[6] Wensveen, F.M., Derks, I.A.M, van Gisbergen, K.P.J.M., de Bruin, A.M., Meijers, J.C.M., Yigittop, H.A., Nolte, M.A., Eldering, E. & van Lier, R.A.W. BH3-only protein Noxa regulates apoptosis in activated B cells and controls high-affinity antibody formation. Blood 119(6), 1440–9 (2012). DOI: 10.1182/blood-2011-09-378877

[7] Zhu, Z., Olson, K. S. & Magliery, T. J. 50 years of antibody numbering schemes: a statistical and structural evaluation reveals key differences and limitations. Antibodies (Basel). Dec 4, 2024. 13(4):99. doi: 10.3390/antib13040099

